# Construction and Characterization of a Synthetic Baculovirus-inducible 39K Promoter

**DOI:** 10.1101/403279

**Authors:** Zhan-Qi Dong, Zhi-Gang Hu, Hai-Qing Li, Ya-Ming Jiang, Ming-Ya Cao, Peng Chen, Cheng Lu, Min-Hui Pan

**Affiliations:** State Key Laboratory of Silkworm Genome Biology, Southwest University, Chongqing 400716, China; Key Laboratory for Sericulture Functional Genomics and Biotechnology of Agricultural Ministry, Southwest University, Chongqing 400716, China; Joint National Laboratory for Antibody Drug Engineering, Institute of Immunology, Henan University School of Medicine, Kaifeng 475004, China.

**Author notes:** Address correspondence to Cheng Lu, and Min-Hui Pan. These authors contributed equally to this work (Zhan-Qi Dong, Zhi-Gang Hu).

**Keywords:** Keyword, *Baculovirus*, *39K*, *Inducible* promoter, *Synthetic*

## Abstract

The low expression activity and specificity of natural promoters limit the applications of genetic engineering. To construct a highly efficient synthetic inducible promoter in the *Bombyx mori* (Lepidoptera), we analyzed the regulatory elements and functional regions of the *B. mori* nucleopolyhedrovirus (BmNPV) 39K promoter. The results of truncated mutation analysis of the 39K promoter showed that the transcriptional regulatory region spanning positions -573 to -274 and +1 to +62 is essential for virus-inducible promoter activity. Further investigation using electrophoretic mobility shift assay (EMSA) revealed that the baculovirus IE-1 protein binds to the 39K promoter at the -310 to -355 region, and transcription activates the expression of 39K promoter assay. Finally, we successfully constructed a synthetic inducible promoter that increase the virus-inducing activity of other promoters using the baculovirus-inducible transcriptional activation region that binds to specific core elements of 39K (i.e., spanning the region -310 to -355). In summary, we describes a novel, synthetic, and highly efficient biological tool, namely, a virus-inducible 39K promoter, which provides endless possibilities for future gene function research, gene therapy, and pest control in genetic engineering.

## Important

Silkworm genetic engineering is widely used in gene function, silk engineering and disease-resistant engineering in most applied in Asia. However, some of the earliest promoter elements are still used to control the development of silkworm transgenic expression and gene therapy. To develop effective genetic engineering technologies for silkworm and baculovirus expression system, we constructed a highly efficiently synthetic baculovirus-inducible 39K promoter in insects. Which successfully constructed and optimized a synthetic inducible promoter 39K that can be effectively applied CRISPR/Cas9 gene editing and transgenic technology to construct transgene material of silkworm and provides an efficient tool for synthetic biology and gene therapy. The synthesized inducible promoters also provides new insights to improve strategies for insect genetic engineering, pest control and gene function research.

## Introduction

The inducible promoter, known as the inducible regulation sequence or the inducible enhancer, is a group of promoters that can enhance the expression of exogenous genes under the stimulation of specific physical, chemical, or pathogen signals (1, 2). In general, the inducible promoter, similar to the transcriptional activator, exists in an inactive form and can be directly or indirectly activated by the corresponding signal. Currently, several technical methods supported by inducible promoters (se.g., Cre-loxp, Tet-On/Tet-Off, and ecdysone and pathogen inducible systems), are widely used in the field of animal and plant genetic engineering, including gene function identification or variety improvements (3-6). Insects are the largest group of organisms on earth. Some insects such as silkworms and bees are of important economic value. However, a highly efficient inducible system that could be extensively used in insect genetic engineering research has not been established to date, and thus it is of utmost significance to construct a pathogenic inducible promoter in disease resistance breeding and gene therapy (7, 8).

The synthetic promoter is a promoter that constructs stronger expression levels by combining a unique combination of different promoter elements and replacing or redesigning sequences with various combinations of promoters (9-11). Previous studies on synthetic promoters in plants have mainly focused on synthetic inducible promoters (12). Synthetic promoters are mainly constructed using cis-regulatory elements that bind to fuse core promoters (13). The construction of different pathogen-inducible promoters could effectively improve the spectrum of transgenic disease resistance in plant disease resistance breeding (12, 14). Alternatively, constructing an inducible promoter in combination with a tissue-specific promoter (e.g., root, stem, leaf) and an inducible promoter contributes to specific tissue-induced expression to improve crop quality, crop robustness, and disease resistance (15). Synthetic promoters have also been reported in animals (11). The construction of these synthetic promoters mainly involves the same direction assembly of different expression control sequences, the application to targeted therapy diseases, andspecific tissue expression of foreign genes (16-18). Synthetic promoters have recently been initiated in insect research, particularly for insect disease breeding.

We previously screened for the *B. mori* nucleopolyhedrovirus (BmNPV)-induced promoter (VP1054, P33, Bm21, Bm122, 39K, P143 and P6.9) activity and found that the 39K promoter had the highest BmNPV-induced transcriptional activity (19). The virus-inducing activity of the BmNPV 39K promoter could be further increased using enhancers such asHr3, Hr5, Polh and PU (19). Simultaneously, the overexpression of an exogenous *hycu-ep32* gene controlled by an inducible 39K promoter shows high antiviral capacity in transgenic lines (20). Furthermore, we constructed a baculovirus-inducible RNA interference system that could inhibits BmNPV replication, is tightly controlled by viral infection, and is not toxic to host cells (21). Moreover, a highly efficient CRISPR/Cas9 gene editing system was constructed with reduced potential off-target effects and high editing efficiency using virus-inducible 39K promoter, which enhances the antiviral ability of *B. mor*i cells (22). Therefore, to improve the efficiency of the virus-inducible 39K promoter for gene function studies, silkworm resistance breeding, and pest control, it is imperative to construct a synthetic promoter in insects.

Therefore, in the present study, we constructed a synthetic inducible promoter by identifying the 39K promoter regulatory regions and binding sites. First, we verified the key functional domains (spanning regions -573 to -274 and +1 to +62) of the 39K promoter by gradually introducing truncating deletions at the 5′ end, 3′ end, and intermediate regions based on the characteristics of the 39K promoter regulatory region as indicated by the dual luciferase report system assay. Then, we constructed a promoter with a shorter promoter sequence and better induction activity by analyzing the regulatory elements of the 39K promoter and associated point mutations. Furthermore, we identified the binding site of baculovirus IE-1 transcriptional activation 39K promoter to the -310 to -355 region. Finally, we analyzed that the 39K promoter-inducing active region combined with specific promoters to construct inducible promoters that can efficiently and specifically activate promoter expression. The results show that we successfully constructed a synthetic inducible promoter 39Kthat can be effectively applied to insect gene function research, disease resistance breeding, and pest control.

## Methods

### Cells and Viruses

The *B. mori* ovary cell line BmN-SWU1 was cultured at 27 °C in TC-100 medium (United States Biological, USA) supplemented with 10% (V/V) fetal bovine serum (FBS) (Gibco, USA) and 10% (V/V) penicillin/streptomycin (23). Recombinant BmNPV (vA4^prm^-EGFP) containing an EGFP marker gene driven by the*B. mori* actin A4 promoter was created from the bacmid bMON7214, which contains the BmNPV genome (21, 24). The BmN-SWU1 cells were transfected with the vA4^prm^-EGFP construct, and viral titers were determined using the 50% tissue culture infective doses (TCID_50_) assay (24).

### Plasmid Construction

Previous studies have shown that the -773 bp upstream and +134 bp downstream motifs of the 39K promoter transcription initiation site are critical regions for 39K promoter activity (19). To analyze the structural features of the 39K promoter, we performed a stepwise truncation analysis of the 39K promoter. Truncated fragments of 39K promoters were cloned into a pGL3-basic vector (Promega, USA) to construct the *Firefly luciferase* (FLUC) expression vector. The 5′ truncated plasmids fragment included P-723 (-773∼-724 deletion), P-673 (-773∼-674 deletion), P-623 (-773∼-624 deletion), P-573 (-773∼-574 deletion), P-523 (-773∼-524 deletion), P-473 (-773∼-474 deletion), P-423 (-773∼-424 deletion), P-373 (-773∼-374 deletion), P-323 (-773∼-324 deletion), P-273 (-773∼-274 deletion), P-223 (-773∼-224 deletion), P-173 (-773∼-174 deletion), P-123 (-773∼-124 deletion), P-73 (-773∼-74 deletion), and P-23 (-773∼-24 deletion) (Figure 1B). The 3′ truncated fragment plasmids included P+116 (+117∼+134 deletion), P+96 (+97∼+134 deletion), P+76 (+77∼+134 deletion), P+62 (+63∼+134 deletion), and P+1 (+2∼+134 deletion) (Figure 1D). The intermediate segment deletion plasmids included ΔP-1∼-223(-1∼-223deletion),ΔP-1∼-273 (-1∼-273 deletion), ΔP-1∼-373 (-1∼-373 deletion),ΔP-1∼-473(-1∼-473deletion),ΔP-223∼-273(-223∼-273deletion),x0394;P-223∼-373(-223∼-373deletion),andΔP-373∼-473(-373∼-473 deletion) (Figure 1C). Then, the sequence of the IE1promoter and the *Renilla luciferase* (RLUC) reporter gene were linked to the pGL3 vector, named as pGL3-IE1-Rluc and used as internal reference plasmid. The baculovirus immediate early genes *ie-0*, *ie-1*, *ie2*, *pe38* and *me53* from the BmNPV genome were cloned into a pIZ/V5-His (Invitrogen, USA) vector to generate pIZ-IE0, pIZ-IE1, pIZ-IE2, pIZ-PE38 and pIZ-ME53. Baculovirus-inducible promoter 39K was cloned into pIZ-DsRed to replace the OpIE2 promoter. The resulting p39K-DsRed plasmid was used as vector backbone for baculovirus-inducible expression of DsRed. All clones were verified by sequencing. All primers used in this study are presented in Table S1.

**Figure 1.**
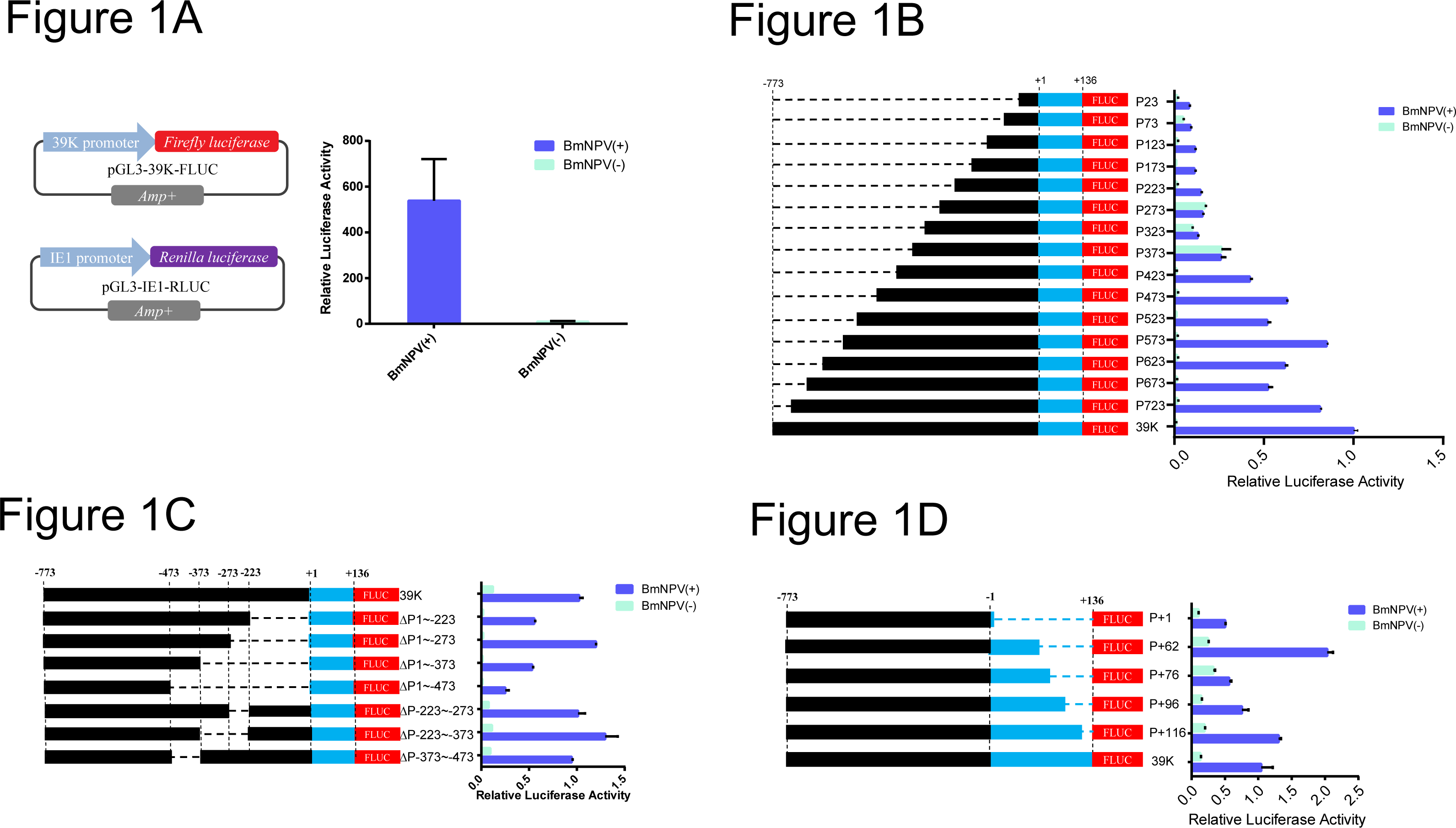
Structural and functional analysis of the 39K promoter. (A) Schematic of *Firefly luciferase* and *Renilla luciferase* expressed vectors to sustain the dual luciferase reporter system. (B) Relative luciferase assay of the 5´-end truncated of the 39K promoter. Cells co-transfected with the *Firely luciferase* and *Renilla luciferase* expression vector were infected or uninfected with BmNPV at 10 MOI. Cells were examined under luciferase reporter system at 48 h p.i.. Black represents -773∼-1 fragment, blue represents +1∼+136 fragment and dashed line represents the missing fragment of the 39K promoter. Red represents the *Firely luciferase* reporter gene. The 39K promoter luciferase activity represents 1 and the promoter activity of the other truncated fragment is a ratio relative to 39K promoter. (C) Relative luciferase assay deletion and truncated fragment of the 39K promoter. (D) Relative luciferase assay 3´-end truncated of the 39K promoter.

### Dual Luciferase Reporter Assays

The dual luciferase expression plasmids pGL3-39K-Fluc (450 ng) and pGL3-IE1-Rluc (50 ng) were co-transfected into the BmN-SWU1 cells. Approximately 24 h later, these were then infected with BmNPV at MOI=10. At 72 hours post infection (h p.i.), the cells were collected, and luciferase activities were measured with using Dual-Glo luciferase Assay kit (Promega) using ultra-high sensitivity fluorescence chemiluminescence detector. Relative luciferase activity (FLUC/RLUC) was normalized to the values obtained using pGL3-39K-Fluc as control plasmid. Each experiment analysis was repeated thrice.

### Transfection and Fluorescence Analysis

The BmN-SWU1 cells were cultured in 24-well plates (Corning, USA). After the cells had stabilized, BmNPV immediate early gene expression plasmids pIZ-IE0, pIZ-IE1, pIZ-IE2, pIZ-PE38 and pIZ-ME53 (0.4 μg) with the p39K-DsRed (0.4 μg) plasmid were co-transfected into the cells using the X-tremeGENE HP DNA Transfection Reagent (Roche, Switzerland). At 48 h post-transfection (h p.t.), all cells were visualized on an Olympus inverted fluorescence microscope with the same parameter settings.

### Reverse Transcription-quantitative PCR (RT-qPCR)

After the BmNPV immediate early gene expression plasmid pIZ-IE0, pIZ-IE1, pIZ-IE2, pIZ-PE38 or pIZ-ME53 with the p39K-DsRed plasmid were co-transfectedinto the cells, total RNA was isolated using the TRIzol RNA extraction kit (ThermoFisher Scientific, USA), following the manufacturer ’s instructions. RT-PCR were performed with an iTaqTM Universal SYBR^®^ Green Supermix and CFX Connect Real-Time PCR Detection System (Bio-Rad, USA) using primers specific for DsRed (Table S1). The *Bombyx mori sw22934* gene was used as the reference. The reaction conditions of RT-PCR were as follows: 95 °C for 30 s; followed by 40 cycles at 95 °C for 5 s and 60 °C for 20 s with 1 M of each primer. All experiments were repeated three times.

### Recombinant Expression and Protein Purification

The coding region of IE-1 was amplified with specific primers IE1-F/IE1-R and cloned into the pCold-I vector and the pGX-4T-1 vector. Positive plasmids were transformed into *E. coli* strain BL21 competent cells and induced with 0.3 mM, 0.5 mM and 1.0 mM, of IPTG to express the IE1-His recombinant protein. The IE1-His protein was purified using a His-Trap HP column (GE Healthcare, Germany), according to the manufacturer’s recommendations.

### Electrophoretic Mobility Shift Assay (EMSA) Analysis

To analyze the potential binding sites of the 39K promoter, two different transcription factor binding site prediction programs, namely, Neural Network Promoter Prediction (http://www.fruitfly.org/seq_tools/nnppHelp.html) and JASPAR CORE (http://jaspar.genereg.net/) were used. A total of four potential transcription factor binding sites were identified, which were located at positions -486 to -532,-386to -431,-310 to -355 and +2 to +47 of the 39K promoter. For EMSA, the probes were5´-labeled with biotin (Thermo Fisher Scientific, USA), and then the labeled oligonucleotides were annealed to produce a double-stranded probe. All probe used in this study are presented in Table S2.

To evaluate the interactions between IE-1 proteins and 39K regulatory elements, EMSA was conducted according to the guidelines of the Light Shift Chemiluminescent EMSA kit (Thermo Fisher Scientific). After a 30 min incubation at 25°C, reaction mixtures were loaded onto 6% (w/v) native polyacrylamide gels and resolved by electrophoresis electrophoresed in TBE buffer (89?mM Tris, 89?mM boricacid, 2?mM EDTA, pH 8.3) for approximately 1 h at 100V on ice. The proteins were transferred onto a PVDF membrane (Roche). Bound HRP-conjugated bands were visualized using the LightShift Chemiluminescent EMSA kit according to the manufacturer’s protocol.

## Construction of the Artificial Inducible 39k Promoter

Based on the results of 39K promoter truncation analysis, three synthetic inducible promoters were constructed, namely, p39K-1 (contains the +1∼+62 and-273∼-573 fragments), p39K-5 (contains the +1 ∼+134 and -273∼-573 fragments) and p39K-9(contains the +1∼+134 and -273∼-773 fragments). To improve the promoter activity of 39K, point mutations of the CAAT box to CGGT at position of -329, -399, or -329 and -399 were created. A total of 12 synthetic inducible promoters were constructed in combination with truncated and point mutation vectors and designated as p39K-1to p39K-12, respectively. All artificially inducible promoters were synthesized by Genscript (Nanjing, China) and cloned into a pGL3-basic vector.

## Statistical analysis

All data were expressed as the mean ± SD of three independent biological experiments. Statistical analyses were performed with student’s *t* tests using GraphPad Prism6. Differences with *P* < 0.01 were considered statistically significant.

## Results

### Structural and Functional Analysis of the 39K Promoter

To generate optimized virus-inducible specific promoters, a truncation and mutation strategy was employed to gradually remove the 39K promoter core region, followed by analysis for change in 39K promoter activity. After the 39K promoter-controlled *Firely luciferase* and the reference plasmid IE1 promoter-controlled *Renilla luciferase* were co-transfected into the BmN-SWU1 cells, promoter activity was assessed by detecting changes in *Firely luciferase* activity relative to that of *Renilla luciferase* (Figure 1A). To identify the core areas required for highly expression, deletion mutants were created. Using -773∼+134 as the original sequence of the 39K promoter, each truncation was reduced by 50 bp relative to the original sequence (Figure 1B). Fifteen 5′-truncated luciferase assay plasmids of the 39K promoter and the reference pGL3-IE1-Rluc plasmid were co-transfected into the BmN-SWU1 cells. At 48 h p.t., the luciferase activity was evaluated by adding BmNPV or the culture medium and incubating for 48 h. The results indicated a gradual decrease in promoter activity with shorter promoter lengths. The length of the P573 promoter was shorter by 200-bp relative to the 39K promoter, but t promoter activity only showed a 14.5% decrease (Figure 1B). Fragment -773∼-573 exhibited little effect on 39K promoter activity. However, the activity of the P-323 promoter decreased by 97.21% relative to the 39K promoter. These findings suggest that the-323∼-573 fragment harbors an important regulatory element of the 39K inducible promoter. The plasmids P273, P323, and P373 showed strong constitutive promoter activity, and that of the P273 promoter was 12.27-fold higher than P223, indicating that the -223-273 fragment was related to the constitutive promoter activity of the 39K promoter (Figure 1B).

To further analyze the 39K promoter regulatory motif, an intermediate deletion fragment of the 39K promoter was created. The results showed that ΔP-1∼-273,ΔP-223∼-273,andΔP-373∼-473 had no significant effect on 39K promoter activity (Figure 1C).FurtherpromoterdeletionfragmentofΔP-1∼-223,ΔP-1∼-373,andΔP-1∼-473, led to a rapid decrease in promoter activity (Figure 1C). Therefore, combined with the 5′-end deletion results and the principle of selecting optimal promoters, the -1 to -273 fragment of 39K promoter could be delete to construction of artificial inducible 39K promoter. The +1∼+134 fragment of the 39K promoter is the core region, and the 3′ end was gradually truncated and the promoter activity was analyzed. The results showed that the promoter activities of P+116 and P+62 increased by 35.4% and 97.00% compared to 39K, respectively. These results indicate that the deletion of +134∼+116 and +76∼+62 increases the activity of the 39K promoter (Figure 1D). These two fragments impart inhibitory effects on promoter activity. Therefore, the optimal promoter would have the +136 to +62 fragments deleted from the 3′ end based.

### Construction of an Artificial Inducible 39K Promoter

Deletion analysis of the 39K promoter identified the regions that have an effect on promoter activity. In addition, we analyzed the key regulatory elements in the core region of the promoter using a promoter prediction program. Online analysis showed that the 39K promoter contains core components such as two enhancer-like components CGTGCGC, six CAAT loci, two transcription inhibitors TGAC, two *cis*-regulatory originals CACT, and two TATA boxes (Figure 2A). In combination with the position of the 39K promoter core element and key regulatory regions, we first constructed three artificial inducible promoters, which included P39K-1 (-573∼-273 and +1∼+62 fragments), P39K-5 (-573∼-273 and +1∼+134), and P39K-9 (-773∼-273 and +1∼+134). The activities of the P39K-1, P39K-5 and P39K-9 promoters were 87.24%, 75.94%, and 112.34% of that of the 39K promoter, respectively (Figure 2B). The promoter lengths of the P39K-1, P39K-5, and P39K-9 promoters were 362 bp, 436 bp, and 636 bp, respectively. The purpose of constructing an artificial promoter was to minimize the length of the promoter without affecting its activity. Therefore, the length of the P39K-1 promoter was only 39.91% of the 39K promoter sequence, but the promoter activity still reached the original 87.24%, which is the better artificially induced promoter. Previous studies have shown that mutations involving of the CAAT site to CGGT significantly increases the promoter activity (25). Therefore, we constructed nine artificial inducible promoters with -326 loci, -399 loci, and two simultaneous mutations in the P39K-1, P39K-5, and P39K-9 promoters, respectively. Dual luciferase reporter assays showed that the promoters of these nine point mutations did not significantly increase promoter activity relative to the P39K-1, P39K-5 and P39K-9 promoters (Figure 2B). The P39K-1 artificially inducible promoter still contains enhancers such as component CGTGCGC, the CAAT locus, and the transcription inhibitor TGAC. After the previous two TATA boxes were deleted, a new TATA box appeared at the -70 bp position of the transcription initiation site of the artificially inducible promoter P39K-1 (Figure 2A). These results indicate that P39K-1 still possesses the original promoter regulatory mechanism and thus an optimized artificial inducible promoter.

**Figure 2.**
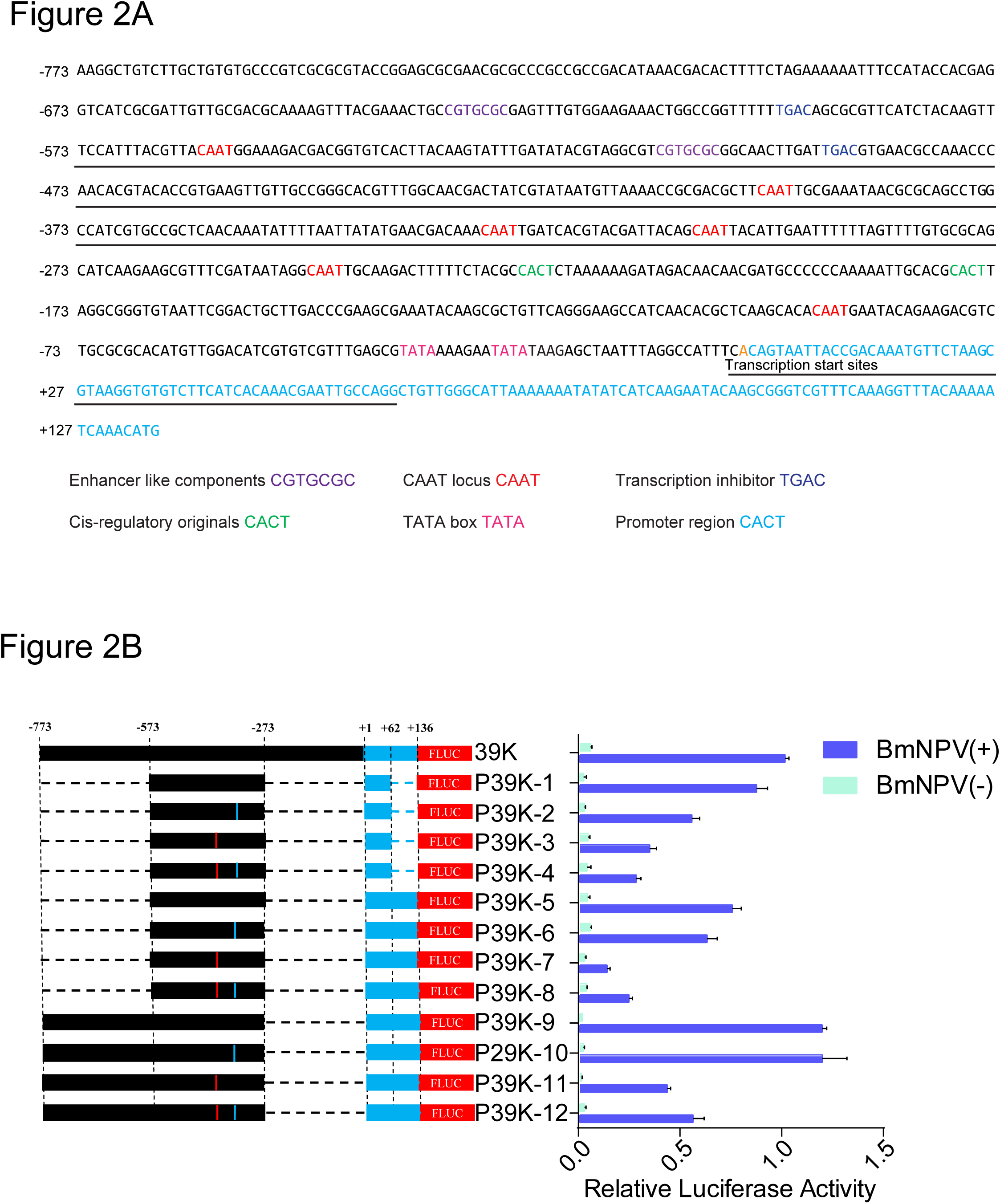
Construction of an artificial inducible 39K promoter. (A) Analysis of 39K promoter regulatory element. Purple represents enhancer like components CGTGCGC element, red represents CAAT locus, blue represents transcription inhibitor TGAC box, green represents cis-regulatory original CACT element, and pink represent TATA boxes. Artificially inducible 39K promoter sequences are underlined. (B) Relative luciferase assay of the artificial inducible 39K promoter. BmN-SWU1 cells were co-transfected with the indicated *Firely luciferase* and *Renilla luciferase* expression vector and infected with BmNPV at 10 MOI or uninfected. At 48 h p.i., cells were examined using a luciferase reporter system. Each data point was determined from the mean of three independent replicates. The red location represents the CAAT mutation to CGGT of 39K promoter -399 site. The blue location represents the CAAT mutation to CGGT of 39K promoter -329 site.

### Identification of Inducible Promoter 39K-regulated Genes

The expression of the baculovirus gene is regulated by the cascade, and subsequent phase gene expression is dependent on the previous phase (26). The baculovirus *39K* gene is a delayed early expression gene (27). To identify the 39K promoter transcriptional control gene, we first screened the transcriptional regulation of the 39K promoter by analyzing five immediate-early genes (i.e., *ie-0*, *ie-1*, *ie-2*, *pe38* and *me53*). We co-transfected pIZ-IE0, pIZ-IE1, pIZ-IE2, pIZ-PE38 and pIZ-ME53 with p39K-DsRed and then detected DsRed at the transcriptional levels. The results indicated the expression of the DsRed protein only in the viral-infected and pIZ-IE1 transfected BmN-SWU1 cells, but not in the pIZ-IE1, pIZ-IE2, pIZ-PE38, pIZ-ME53, and non-infected cells (Figure 3A). These results indicate that the DsRed protein is rapidly activated by viral infection and IE-1 protein expression. Moreover, to detect the sensitivity of the inducible promoter, we investigated the transcription of DsRed as induced by viral protein and BmNPV. The results showed that the virus and IE-1 protein induced large-scale transcription of *DsRed* (Figure 3B). No changes in *DsRed* transcription levels in the pIZ-IE0, pIZ-IE2, pIZ-PE38, pIZ-ME53 transfected and non-infected cells were observed. The luciferase assay also showed that only the IE-1 protein could effectively induce 39K promoter activity. In addition, the other early genes were not transcriptionally regulated by the 39K promoter.

**Figure 3.**
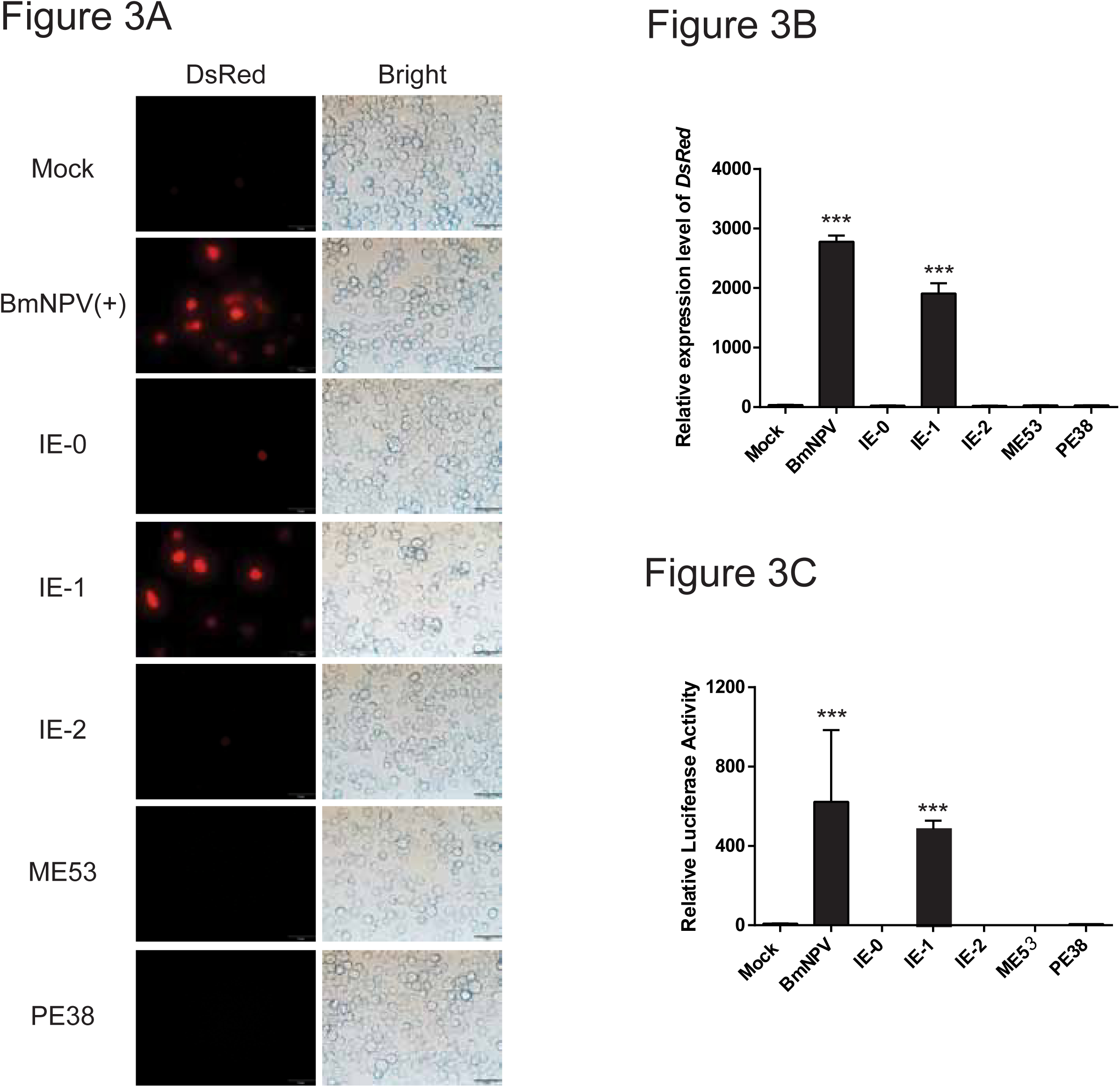
Identification of inducible promoter 39K-regulated genes. (A) Immunofluorescence analysis of 39K promoter activated foreign protein expression. p39K-DsRed plasmid co-transfection with immediate early genes and examined under a fluorescence microscope at 96 h p.i. Red represents DsRed protein expression, white represents the number of cells. (B) Transcription of inducible p39K-DsRed systemwith BmNPV immediate early genes. Transient co-expression of p39K-DsRed plasmid and immediate early gene or infected with BmNPV at 10 MOI. At 48 h p.i., total RNA was isolated from each transfected cell and quantified by RT-PCR. Each data point was determined from the mean of three independent replicates. (C) Relative luciferase assay of inducible p39K-DsRed system with BmNPV immediate early genes. Each data point was determined from the mean of three independent replicates. ** represent statistically significant differences at the level of *P* < 0.01.

### EMSA Analysis of IE-1 Binding to the 39K Promoter Region

To further strengthen the argument that IE-1 is a direct transcriptional target of the 39K promoter, we performed a gel-shift competition assay using a biotin-labeled oligonucleotide spanning the potential IE1-binding sequence as probe. Through online program prediction, we designed a total of four probes containing multiple potential binding sites. These probes were named probe 1 (-486∼-532), probe 2 (-386∼-431), probe 3 (-310∼-355), and probe 4 (+2∼+47), which were incubated with purified IE-1 derived from prokaryotic expression. The incubation of the biotin with the IE-1 protein. The incubation of biotin labelled probe 3 (-310∼-355) with IE-1 protein resulted in a distinct band shift in the EMSA, which disappeared with the addition of competitive unlabeled DNA probes (Figure 4A). In contrast, no significant band shift was detected in the EMSA after incubation with probe 1 (-486∼-532), probe 2 (-386∼-431) and probe 4 (+2∼+47) (Figure 4A).

**Figure 4.**
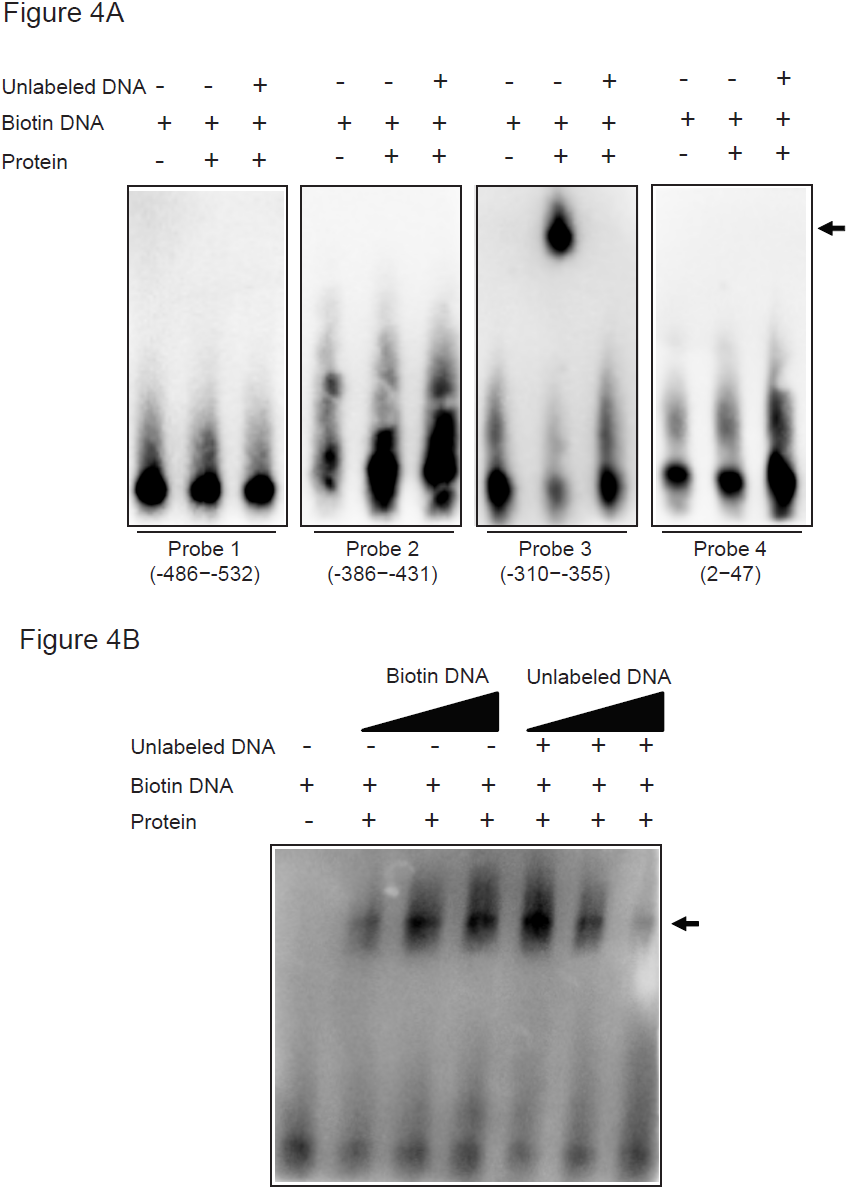
EMSA analysis of IE-1 binding to 39K promoter region. (A) Electrophoretic mobility shift assay indicated that the 39K probes bind to the recombinant IE-1 proteins. We used the competitive inhibitors unlabeled DNA probes as control and without IE-1 proteins as negative control. The shift of the positive control is indicated by a thick stripe. We detected probes 3 (-310∼-355) with block stripes. On the contrary, probe 1 (-486∼-532), probe 2 (-386∼-431), and probe 4 (+2∼+47) were not associated with IE-1. (B) The EMAS detected that the probe 3 (-310∼-355) binds to the recombinant IE-1 proteins. The probe 3 (-310∼-355) probes concentrations were 1, 2, and 6 pmol/L; the IE-1 protein concentration was 0.8 μg/L; the concentrations of compete probes were 2, 20, and 100 pmol/L.

To further examine the binding activity of probe 3 with the IE-1 proteins, we analyzed the effect of the biotin labelled probe and unlabeled DNA on band shifting. The results showed that the incubation of probe 3 with the IE-1 protein resulted in a band shift, which increased with greater biotin labelled probe 3 concentrations and reduced with increasing concentrations of competitive unlabeled DNA probes (Figure 4B). No significant band shift was detected in the probe without incubation with the IE-1 proteins. There results indicate indicating that IE-1 specifically binds to 39K promoter probe 3 (-310 to -355) during the transcriptional activation of the BmNPV IE-1 protein-inducible 39K promoter.

### Application of Artificial Inducible 39K Promoter

To expand the potential use of artificially-inducible 39K promoters in insect genetic engineering, we synthesized new promoters P33+39K(-310∼-355) by combining baculovirus P33 promoters and 39K(-310∼-355) binding sequences. Dual luciferase assays indicated that the P33+39K(-310∼-355) promoters exhibited a significant increase in activity compared to the original sequence after binding to the 39K sequence (Figure 5A). The promoter activity of P33+39K(-310∼-355) increased by4.46 fold after viral infection, which was 1.48-fold higher than the original sequence (Figure 5A). These results demonstrate that the 39K promoter fragment can be utilized in the construction of an artificially inducible promoter to increase induction activity in genetic engineering.

**Figure 5.**
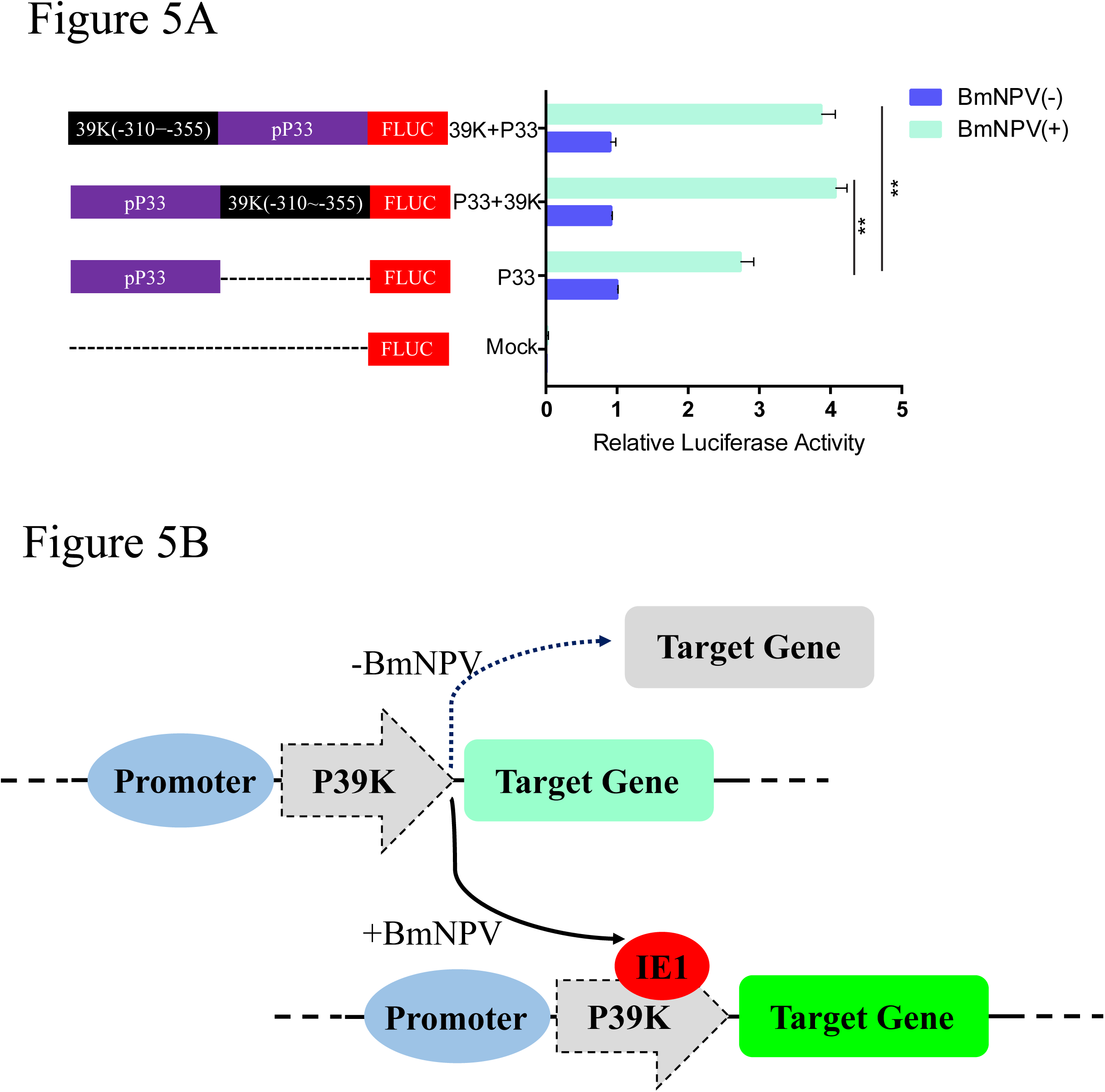
Application of the artificial inducible 39K promoter. (A) Relative luciferase assay of the artificial inducible 39K promoter. BmN-SWU1 cells were co-transfected with indicated *Firely* luciferase and *Renilla* luciferase expression vector and infected with BmNPV at 10 MOI or uninfected. At 48 h p.i., cells were examined under luciferase reporter system. Each data point was determined from the mean of three independent replicates. NS, not significant. ** represent statistically significant differences at the level of *P*< 0.01. (C) Schematic plot of the synthetic inducible promoter project.

## Discussion

Naturally occurring promoters are currently used in protein production and gene therapy (28, 29). However, natural promoters are not always capable of driving high levels of gene expression and may also lack the required specificity depending on the promoter and the specific application (28, 30). As genetic engineering goals become more elaborate and targeted, more precise gene expression tools will be needed (31, 32). Synthetic promoters contain fragments of natural promoters to form new DNA sequence fragments that are not found in nature and are more powerful and specific than naturally occurring promoters (11, 32). Considering that scientists have been engineering silkworms for more than 20 years and that silkworm genetic engineering has been widely used in gene function, silk engineering, and disease resistance breeding in most applied in Asia, it is surprising that we are still using some of the earliest-developed tools to control transgene expression in silkworms (33-36). To more effectively and specifically apply silkworm genetic engineering, we constructed a highly efficient synthetic baculovirus-inducible39K promoter. The 39K (-310∼-355) sequence widely used to enhance other promoter activities to construct synthetic inducible promoters provides an efficient tool for synthetic biology and genetic engineering.

In our previous studies, we have shown that the P-44 (-44 to +133) and (from-420 to -611) are important regions for the transcriptional activation of the 39K promoter, although the activity of the 39K promoter induced by the transcriptional regulatory region was not studied in detail (19). To obtain a synthetically inducible promoter with a shorter sequence and better induction activity, we performed a stepwise analysis of the 39K promoter transcriptional regulatory region to identify the influence of different regions on 39K promoter and induction activity. In combination with the above promoter activity analysis, we constructed three artificially inducible promoters P39K-1 (-573∼-273 and +1∼+62 fragments), P39K-1 (-573∼-273 and+1∼+134), and P39K-9 (-773∼-273 and +1∼+134). Previous studies have shown that mutation of the *Autographa californica* multiple nucleopolyhedrovirus (AcMNPV) ubiquitin promoter CAAT into CGGT increases inducible promoter activity (25). Inthe present study, we compared mutations in the CAAT site of each artificially inducible promoter, which did not exhibit a significant increase compared to the original promoter (Figure 2). Therefore, the optimal synthetic inducible promoter for P39K-1 was constructed in this study. The promoter length of P39K-1 was only 33% of the P39K promoter, without a significant decrease in promoter activity relative to the P39K promoter. The reduction of such long promoter fragments provides significant improvement to the field of genetic engineering.

The AcMNPV 39K promoter is mainly expressed by immediate early genes (37-39). To systematically analyze transcriptional activation of BmNPV, we analyzed the expression of the 39K promoter as induced by different early transcriptional activator factors. The results showed that only the IE-1 could induce 39K promoter initiation activity, but that of IE-0 and IE-2 did not, unlike the AcMNPV 39K promoter (Figure 3) (27, 38, 39). The promoter-specific application could be significantly improved by the expression of the promoter according to the inducible promoter regulatory sequence binding specificity in plant genetic engineering and mammalian gene therapy applications (6, 18, 32). Here, we demonstrate that the 39K (-310∼-355) sequence can be applied to the construction of artificially inducible promoters (Figure 5B). Furthermore, we also could use the original sequence to increase the induction activity of other weakly expressed promoters or increase the inducible activity of a promoter by repeating this fragment several times. Meanwhile, the combination of different promoter regulatory elements also could be used to improve the activity of synthetic inducible promoters. In our previous studies, we successfully applied the virus-inducible promoter 39K to transgenic overexpressing foreign genes, RNAi, and gene editing, and the determination of this promoter binding region may be more accurately applied to the regulation of genetic engineering (20-22). These synthetic inducible promoters also allows more extensive applications to biopharmaceutical and agricultural process and in novel gene therapies. The successful construction of baculovirus synthetic inducible promoters provides a new strategy for the research and application of insect genetic engineering, pest control, baculovirus expression systems, and insect bioreactors. In our future research, we plan to use the following strategies to improve the scope of applications of virus-inducible promoters: 1) incorporating inducible promoter regulatory sequences and tissue-specific promoters to synthesize new promoters to induce expression of specific proteins in specific tissues to avoid loss of host energy and cell cytotoxicity. 2) combined with 39K promoter and IE1 protein binding sequence, a foreign protein inducible expression system will be constructed and applied to insect gene function research; and 3) a broad-spectrum pathogen induction system will be constructed to cultivate genetic engineering varieties that could respond to different pathogens. In addition, the optimization of synthesized promoters can further increase the specificity and yield of foreign proteins expressed by baculovirus expression systems, as well as the application of insect pest control, such as pathogen-inducible transgenic cotton bollworm, *Spodoptera exigua*, and the other economic crop pests. In conclusion, the successful construction of synthesized inducible promoters provides new insights to improve strategies for insect genetic engineering, pest control and gene function research.

## Author contributions

Z.D. and Z.H. performed the vector cloning, sequencing, cell cultures and PCR. Z.D., Y.J. and Z.H. performed the protein purification and EMSA analysis. Y.J., Z.H., andM.C. performed the qRT-PCR and dual luciferase reporter assays. Z.D., M.P., andC.L. conceived the experimental design and helped with date analysis. Z.D., M.P., P.C., and C.L. preparation of the manuscript. The final manuscript was reviewed and approved by all authors.

## Figure legends

**Supplemental Table 1. Sequences of primers used in this study.**

**Supplemental Table 2. Sequences of probes used in this study.**

